# Antidiabetic effects of astaxanthin partly mediated by the inhibition of carbohydrate digestion and absorption and *in vitro* DPPH and lowering lipids in high-fat-fed albino mice

**DOI:** 10.1101/2025.06.24.661370

**Authors:** MD Sifat Rahman, JMA Hannan, Jaber Ahmed Shihab, Prawej Ansari

**Affiliations:** Department of Pharmacy, School of Pharmacy and Public Health, Independent University, Bangladesh (IUB), Dhaka 1229, Bangladesh; Department of Pharmacology, National Medical College and Teaching Hospital, Parsa, Birgunj 44300, Nepal

**Keywords:** Astaxanthin, Type 2 diabetes mellitus, Antihyperglycemic agent, Starch digestion, Glucose diffusion, HFF mice

## Abstract

Astaxanthin (ASX) has been reported to reduce hyperglycemia by improving insulin secretion and insulin resistance. The present research aims to elucidate the antidiabetic effects as well as the mode of action of ASX, both *in-vitro* and *in-vivo*. Starch digestion, glucose diffusion, 2,2-Diphenyl-1-picrylhydrazyl (DPPH) were investigated *in vitro,* and oral glucose tolerance tests (OGTT), total residual unabsorbed sucrose contents in the gut, gastrointestinal (GI) motility were assessed in high-fat-fed (HFF) hyperlipidemic albino mice. Acute and chronic metabolic effects were evaluated by monitoring food and water intake, amount of urination and defecation. HFF-induced type 2 diabetic mice were treated for 33 days with ASX (30 and 60 mg/5 ml/kg b.w.) were used to assess chronic effects.

In the *in vitro* study, ASX (1.6 - 1000 µg/mL and 40 - 5,000 µg/ml) decreased (*p<*0.001) enzymatic digestion of starch and glucose diffusion by 45.14% and 32.50%, respectively and also showed a significant inhibitory effect on the DPPH (*p<*0.001) over a concentration range (1.6 - 5000 µg/mL) of ASX. During the *in vivo* experiments, oral administration of ASX (30 - 60 mg/5 ml/kg b.w.) improved oral glucose tolerance (*p<*0.001) as well as significantly increased (*p<*0.01) residual unabsorbed sucrose contents in the six segments of the gut. In addition, ASX (30 - 60 mg/5 ml/kg b.w.) significantly enhanced GI motility (p<0.05) in albino mice. Furthermore, ASX (60 mg/5 ml/kg b.w.) reduced food and fluid intake, and the amount of urine and stool formation significantly (*p<*0.001) both in chronic and acute metabolic studies, accordingly. In a chronic study, ASX, at (30 and 60 mg/5 ml/kg b.w.), substantially lowered blood glucose, total cholesterol, VLDL, triglycerides, LDL cholesterol and also improved liver glycogen content.

In conclusion, ASX shows effect on decreasing carbohydrate digestion, absorption, increasing GI motility, and inhibiting DPPH. Thus, ASX might be a potential drug for the management of diabetes and its complications.

## 1. Introduction

Diabetes mellitus, a long-term metabolic disorder, comprises a complex pathophysiology leading to in prolonged organ damage that results from hypoglycemia and hyperglycemia [1], [2]. The worldwide prevalence of diabetes mellitus in adults in the age 20 to 79 years rises above 10.5%, which affect over 530 million individuals [3]. Global predictions suggest that this number will exceed 643 million in 2030 and nearly 783 million in 2045.[4]. Type 1 Diabetes Mellitus (T1DM), Type 2 Diabetes Mellitus (T2DM), Maturity Onset Diabetes of the Young (MODY), gestational diabetes, neonatal diabetes, and secondary diabetes caused by hormone-related disorders are different types of Diabetes Mellitus (DM).[5]. Diabetic complications comprise microvascular diseases includes nephropathy, retinopathy, and neuropathy, as well as macrovascular complications such as cardiovascular disease, stroke, and arterial disease. Chronic diabetes leads to severe life-threatening complications such as hyperglycemia caused by ketoacidosis [6]. Diabetes, an alarming worldwide health concern, is managed with anti-diabetic medication such as insulin, metformin, sulfonylureas, and different oral agents, illustrating several types of treatment options [7], [8]. Although the therapeutic benefits in managing diabetes mellitus, frequently used medications for DM result in numerous side effects, such as hypoglycemia, fluid retention, osteoporosis, and heart failure, which have been observed resulting from the consumption of oral anti-diabetic treatments [9] Despite their side effects, it is necessary to discover a low-toxicity and efficacious functional food or medication for both the prevention and management of diabetes [10].

Carotenoids, natural phytochemicals present in plenty of consumable plants and fruits, have already shown the ability to modulate immune-inflammatory reactions and regulate non-cytokine intermediaries, including reactive oxygen molecules and nitric oxide, therefore maybe preventing regarding complications associated with diabetes [11]. Several numbers of research have emphasis on the protective effects of carotenoids to prevent acute signs of T2DM [12], [13], [14]. In recent times, considerable focus has been directed towards investigating the hypoglycemic mechanisms associated with dietary carotenoids, which have shown effects that expand beyond mere antioxidant properties [15].

ASX is a xanthophyll carotenoid that contains two terminal rings linked by a polyene chain [16], [17], commonly found in seafood and is mostly produced by microalgae *Haematococcus pluvialis* [17] ASX is known for its medicinal properties, including anticancer, anti-ulcerative, and anti-inflammatory activities [18] and antidiabetic activities [19]. Notably, Numerous reports have highlighted the important role of astaxanthin in diabetes prevention. For instance, astaxanthin had a notable anti-hyperglycemic impact in type 2 diabetic mice [20]. Previous studies also report that ASX could reduce the adverse effects of oxidative stress caused by hyperglycemia in pancreatic β-cells, resulting in enhanced glycemic index and elevated insulin levels in the bloodstream [21]. In addition, a number of studies shown that administration of ASX improved insulin sensitivity in spontaneously hypertensive, obese rats and mice that consumed high-fat and high-fructose diets [20], [22]. Moreover, ASX has been reported in several studies to protect diabetic nephropathy by the suppression of oxidative damage and the mitigation of renal inflammation [23], [24], [25].

Although the antidiabetic effects of ASX is well established, the specific mechanisms through which ASX exerts its antidiabetic effects are still unknown, and information about its impact on carbohydrate digestion and absorption, gastrointestinal (GI) motility, and radical scavenging activity are lacking. This research aims to explore the inhibitory effects of ASX on carbohydrate digestion and absorption, glucose regulation, evaluation of liver glycogen content and lipid profiles and its free radical scavenging activity through both *in-vitro* and *in-vivo* experiments. Thus, these findings explore whether ASX could be used as an adjunct therapy for the management of diabetes and its complications.

## 2. Materials and method

### 2.1. Astaxanthin

The R&D grade Astaxanthin ≥97% (HPLC), isolated from Blakeslea trispora and were procured from (SIGMA-ALDRICH, Co 3050 Spruce Street St. Louis, MO 63103).

### 2.2. In vitro starch digestion assay

Initially, 100 mg of starch (Sigma-Aldrich, St. Louis, MO, USA) was solubilized in 3 milliliters of distilled water, either containing or without the addition of ASX (SIGMA-ALDRICH, Co 3050 Spruce Street St. Louis, MO 63103) at doses between 0.32 to 1000 µg/ml, and the glucosidase inhibitor acarbose at the same concentration range, which was used as a reference standard. The solution was mixed with 40 µl of thermally resistant α-amylase that was produced by Bacillus licheniformis (Sigma-Aldrich, St. Louis, MO, USA). It was then maintained at 80°C for 20 minutes before the mixture was made up to 10 milliliters. A sodium acetate buffer (0.1 M) solution (pH 4.75) and 0.1% amyloglucosidase (30 µL), originating from Rhizopus mold (Sigma-Aldrich, St. Louis, MO, USA), were introduced into 1 milliliters of the preparation. The final formulation was incubated for 30 minutes at 60°C. subsequently the addition of the GOD-PAP (glucose oxidase/four-aminophenazone-phenol) reagent, glucose release from the sample was measured using an ELISA microplate reader [26].

### 2.3. In vitro glucose diffusion test

The glucose diffusion model in vitro employed a cellulose ester (CE) dialysis tube with dimensions of 20 centimeters by 7.5 millimeters, involving a Spectra/Por® CE membrane layer with a molecular weight end (MWCO) of 2000, manufactured by Spectrum in Amsterdam, Netherlands. The dialysis tube was filled with 220 mM glucose and a 2 milliliters formulation of 0.9% sodium chloride solution (BDH Chemicals Ltd., Poole, UK), either with or without ASX (40–5000 mg/mL). This particular agent is recognized for its ability to enhance viscosity and decrease the rate of glucose absorption in the intestines. Each tube was sealed at both ends and put into 50 milliliters centrifuge vessels (Orange Scientific, Orange, CA, USA) holding forty-five milliliters of 0.9% sodium chloride. The vessels were positioned on a rotary shaker and kept at 37 °C. Samples were collected for glucose analysis as previously described. [27].

### 2.4. In vitro DPPH assay

The ability of ASX to scavenge free radicals was analyzed utilizing the 2,2-Diphenyl-1-picrylhydrazyl (DPPH) assessment. For the preparation of 1 mL of the solution, a range of concentrations of ASX and L-ascorbic acid, from 1.6 to 5000 µg/mL, were utilized. Subsequently, 2 mL of a 0.2 mmol/L DPPH solution has been added and mixed thoroughly with methanol using a vortex mixer. The control mixture was produced by agitating 2 milliliters of 0.2 mmol per Liter DPPH with 1 milliliter of distilled water. All mixes were kept in darkness for 30 minutes. Following the specified period, a UV spectrophotometer was employed to determine the absorbance of solutions at 517 nm [28].

### 2.5. Animals, induction of diabetes and study design

Albino mice were procured from International Center for Diarrheal Disease Research, Bangladesh (ICDDRB). The mice were exposed to a 12-hour light and dark period at the controlled temperatures of 22.5 °C and 55-65% humidity at IUB animal house. Fresh water and the standard rat pellet diet were available *ad-libitum* during the entire experiment. The standard diet was composed 38.5% fiber, 36.2% carbohydrates, 20.9% protein, and 4.4% fat [29]. Prior to commencing the tests, the animals were fed a high-fat diet over 6–8 weeks, consisting of 20% protein, 45% fat, and 35% carbs, yielding a nutritional value of 26.15 kJ/g and by fourteen weeks of age, diabetic symptoms had been noticed [30]. Then, at the age of 12 weeks, the mice underwent glucose response to an oral load (OGTT; 2.5g/kg, b.w.). Following that, the mice were categorized as having T2DM if their blood glucose range were between 8 and 12 mmol/L recorded [31].

The groups were divided as follows:

Group 1: Normal control

Group 2: HFF diet control

Group 3: HFF diet control + astaxanthin (30 mg/5 ml/kg)

Group 4: HFF diet control + astaxanthin (60 mg/5 ml/kg)

Group 5: HFF diet control + metformin (50 mg/5 ml/kg)

### 2.6. Acute Metabolic study

Metabolic cages were utilized to monitor food and fluid consumption, fecal quantity, and urine volumes in type 2 diabetic mice that were induced with HFF. Before the experiment, the mice were kept to a 12-hour fasting interval followed by a 24-hour adaptation stage. The diabetes control group received only saline, whereas the treatment groups were administered ASX at dosages of (30 and 60 mg/5 ml/kg b.w.). The positive control group was given metformin at a dosage of (50 mg/5 ml/kg b.w.). The dietary and liquid consumption of every single group was carefully monitored, along with the quantity of stool produced and the volume of urine excreted. The four variables were evaluated hourly during the initial six hours, subsequently at two-hour intervals for the next six hours, and finally following a 24-hour break [32].

### 2.7. Acute oral glucose tolerance test

The impacts of ASX on acute oral glucose tolerance at 0, 15, and 30 days were tested in 12-hour-fasted HFF-induced diabetic mice. Blood specimens were acquired from from the tail end at 0, 30, 60, 120, and 180 minutes, following oral administration of ASX at doses of (30 and 60 mg/5 ml/kg b.w.). The positive control group received metformin at (50 mg/5 ml/kg b.w.) along with glucose at a concentration of (18 mmol/kg b.w.). Blood glucose levels were subsequently assessed using control and negative control groups as previously described [33].

### 2.8. Chronic study of astaxanthin on blood glucose, body weight, food, and fluid intake

The long-term effects of ASX (30 and 60 mg/5 ml/kg b.w.) were evaluated by administering ASX twice a day to diabetic mice over a period of 33 days. The control group of mice received only saline, whereas the positive control group was administered metformin at (50 mg/5 ml/kg b.w.). Assessments of body weight, food and fluid intake, and blood glucose levels, obtained from the tail-tip, were conducted from day 0 to day 33 using ACCU-FAST blood glucose test strips at 3-day intervals [34].

### 2.9. Gastrointestinal motility test

To assess the effect of astaxanthin on gastrointestinal motility, barium sulfate milk was administered. 10% BaSO4 was combined with 0.5% carboxymethyl cellulose to prepare the BaSO4 milk. ASX (30 and 60 mg/5 ml/kg b.w.) and the positive control group, metformin (50 mg/5 mL/kg b.w.), were given to 12-hour fasted HFF-induced diabetic mice 60 minutes prior to the oral gavage of the BaSO4 milk. The control group was administered saline at an oral dose of 10 ml/kg. Both the control and BaSO4-treated mice were sacrificed fifteen minutes later. Representing as a (%) of the overall intestinal diameter, the portion of the small intestine that the BaSO4 milk had traversed between the ileocecal junction and the pylorus was shown [35].

### 2.10. Unabsorbed sucrose content in the gut

The impact of ASX on gut sucrose content was assessed by quantifying the remaining sucrose levels in the gastrointestinal tract after a load of oral sucrose. An oral dose of 50% sucrose solution (2.5 g/kg b.w.) was given to rats with type 2 diabetes after they had fasted for 12 hours and received the ASX (30 and 60 mg/5 ml/kg b.w.), respectively. Blood specimens were obtained from a tail vein at the beginning point 0 minutes and at 30, 60, 120, and 240 minutes following oral treatment to evaluate glycemic levels. To analyze the remaining sucrose accumulation in the gastrointestinal tract, a subset of rats was killed at every time interval. The gastrointestinal system was excised and dissected into the stomach, small intestine, caecum, large intestine, and the small intestine was further divided into upper, middle, and lower 20 cm sections. Each segment underwent rinsing with pre-cooled, pH-adjusted saline and was subsequently centrifugation at 3000 rpm (equivalent to 1000×g) for a duration of 10 minutes. The supernatant underwent boiling for 2 hours to break down residual sucrose, subsequently neutralized using sodium hydroxide. The concentration of glucose in the blood and the quantity of glucose released were quantified from the unabsorbed sucrose remaining in the gastrointestinal tract. GI tract sucrose levels was assessed by measuring the glucose released from hydrolyzed sucrose.

### 2.11. Liver glycogen content measurement

The liver glycogen content was assessed with the anthrone technique [36]. The liver was quantified and mixed in ten milliliters of a 5% trichloroacetic acid (TCA) solution. Following the filtration process to filter out precipitated proteins, a one milliliter volume of the clear supernatant was combined with 2 milliliters of 10N KOH and incubated at 100 °C for one hour. After cooling, 1 mL of concentrated acetic acid solution was introduced, and the mixture was brought up to 10 milliliters introducing deionized water. 1 mL of the following solution was combined on ice with 2 milliliters of a prepared anthrone mixture (100 mg of anthrone was prepared in 50 mL concentrated H_₂_SO_₄_), boiled for 10 minutes, and subsequently cooled. The absorbance at 490 nm was determined with a microplate spectrophotometry [37].

### 2.12. Lipid Profile measurement

For 33 days, HFF induced type 2 diabetic mice received oral therapy twice a day with ASX (30 and 60 mg/5 mL/kg b.w.) or metformin (50 mg/5 mL/kg b.w.), and during this time, lipid levels were evaluated as already outlined [38]. Blood specimens were collected from the tail end into heparinized microcentrifuge vessels (Sarstedt, Numbrecht, Germany) to inhibit coagulation and subsequently centrifugation at 12,000 rpm over a period of five minutes. Plasma was subsequently isolated, and low-density lipoprotein (LDL), triglyceride (TG), very low-density lipoprotein (VLDL) and total cholesterol (TC) contents were assessed utilizing COD-PAP and GPO-PAP (Elabscience Biotechnology Co., Ltd., TX, USA) kits in an automatic tester [39].

### 2.13. Statistical analysis

GraphPad Prism 5.0 for Windows was used to perform the statistical analyses. The results are presented as mean ± SEM. Unpaired Student’s *t* test and one-way and two-way ANOVA with Bonferroni post hoc tests (where applicable) were used to analyze the results. P-values less than 0.05 was regarded as statistically significant.

## 3. Results

### 3.1. Effects of Astaxanthin on in vitro starch digestion

ASX at (1.6 - 1000 µg/mL) exhibited substantial reduction in enzymatic glucose digestion of starch by 11.40% to 45.14% (*p<*0.05 and *p<*0.001; Figure 1B) concentration-dependent inhibition in contrast to the control group. Whereas the Acarbose as a positive control (1.6 - 1000 µg/mL) produced by 28.96% to 81.12% (*p<*0.001; Figure 1A) concentration-dependent inhibition on glucose liberation from starch.

**Figure 1:**
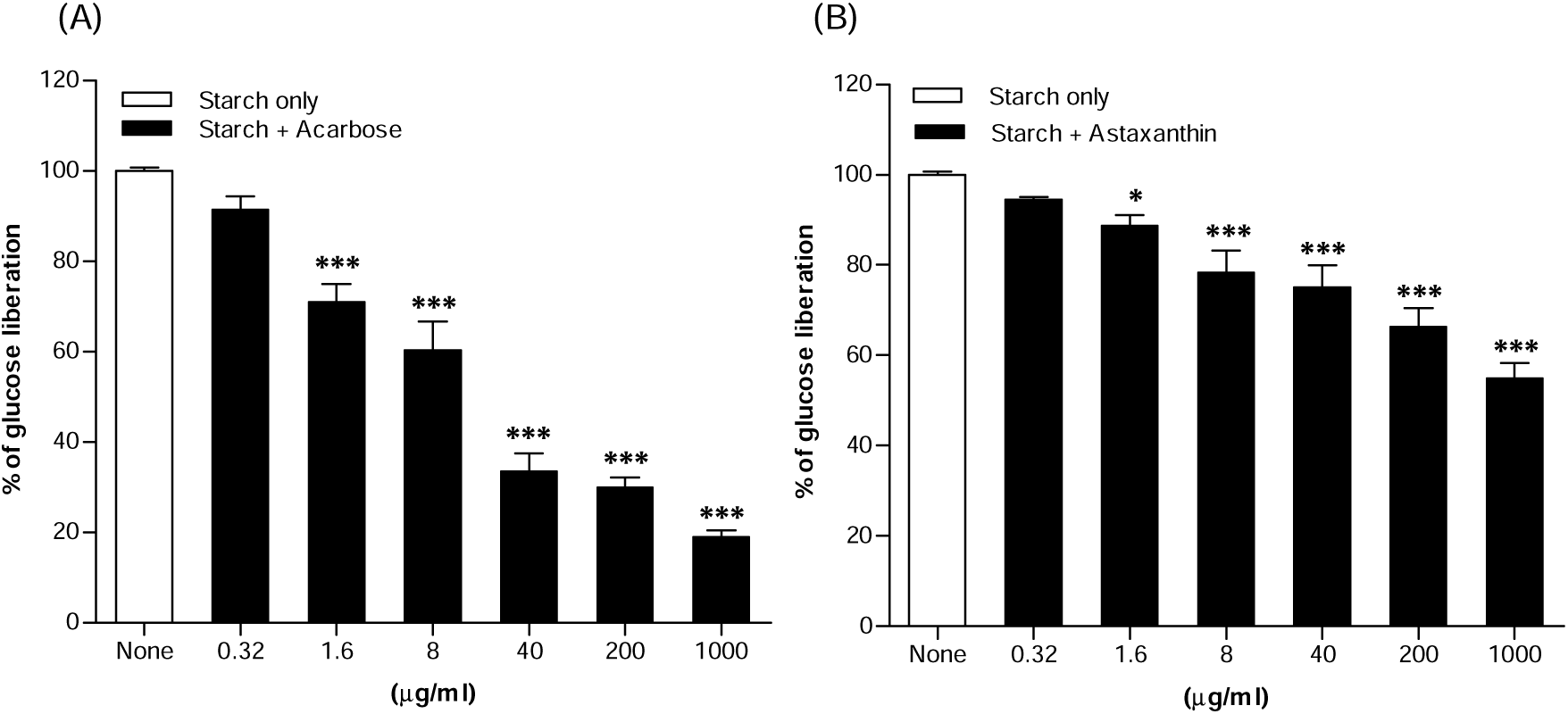
Dose-responsive impacts of Astaxanthin on *in vitro* starch digestion; (A) Acarbose,(B) Astaxanthin. The study was conducted in the including and excluding of positive control acarbose (0.32 - 1000 µg/ml) and ASX (0.32 - 1000 µg/ml) followed by incubation with 0.01% α-amylase and 0.1% amyloglucosidase. Results (n = 4) are expressed as mean ± SEM. **\****P<*0.05**, \*\****P<*0.01**, \*\*\****P<*0.001 comparison to control.

### 3.2. Effects of Astaxanthin on in vitro glucose diffusion

The *in vitro* impacts of ASX on glucose diffusion were assessed corresponding to incubation times of 0, 3, 6, 12, and 24 hours. ASX markedly diminished glucose diffusion and absorption occurred in a manner dependent on both dose and time, across doses from 40 to 5,000 µg/ml. No notable reduction of glucose permeability was detected at 0 hours with ASX (Figure 2A). A notable decline (*p<* 0.05 – 0.01) was detected at 3, 6, and 12 hours in a dose-dependent manner, ranging from 9.83% to 16.63%, 17.87% to 27.66%, and 17.87% to 29.81% at doses of 200 to 5000 µg/mL (Figure 2B, C, D). At 24 hours, the most substantial reduction in glucose absorption was noted, ranging from 15.64% to 32.50% at concentrations 40 – 5000 µg/mL (*p<*0.01 - 0.001; Figure 2E).

**Figure 2:**
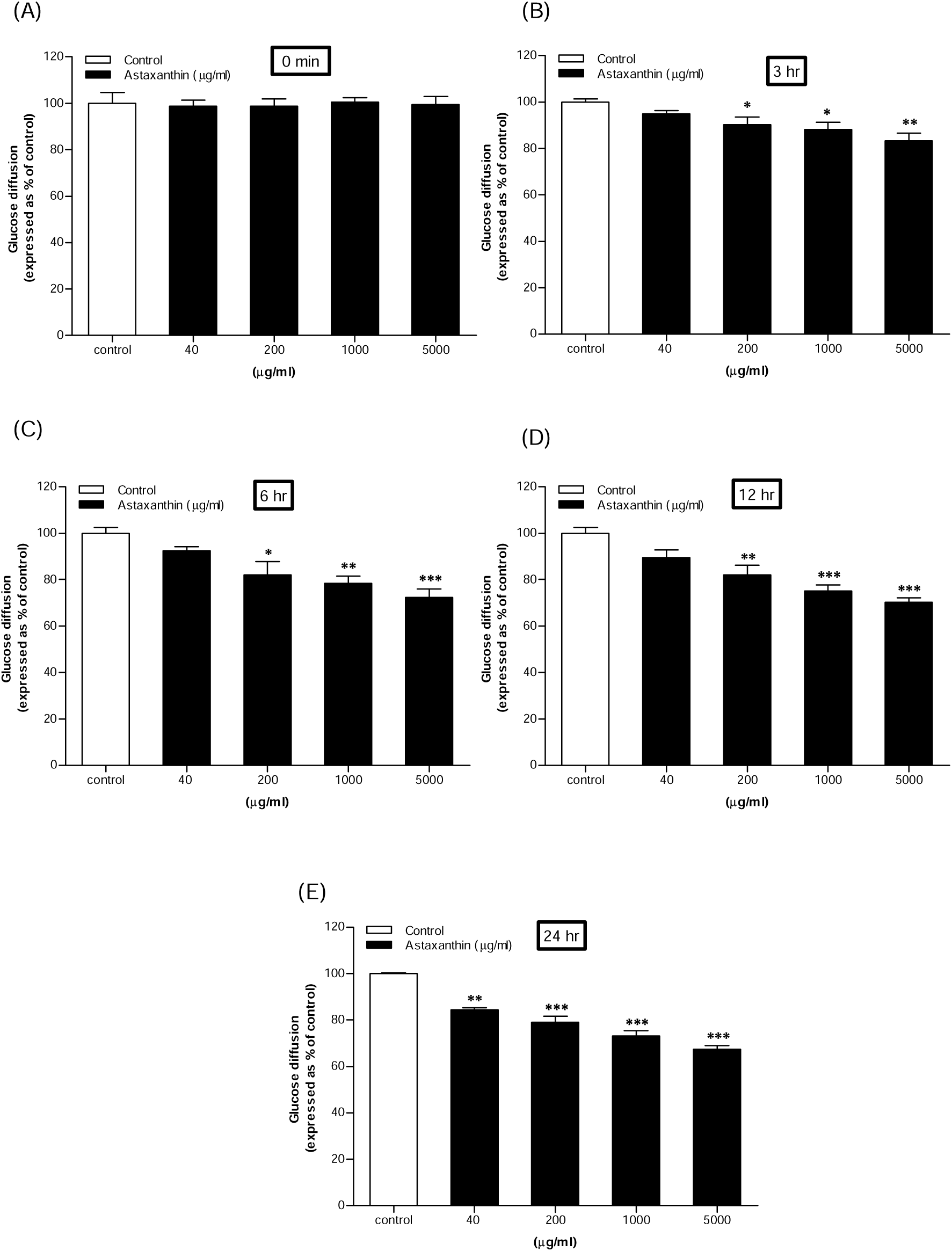
Dose-responsive impacts of Astaxanthin (A, B, C, D and E) on glucose diffusion *in vitro*. The study was carried out in the presence or absence of ASX (40 - 5000 µg/ml) utilizing dialysis tubing made of cellulose membrane, and monitoring the glucose diffusion at (A) 0, (B) 3, (C) 6, (D)12 and (E) 24 hours respectively are depicted as bar graphs, subsequent to the GOD-PAP technique. Results (n = 4) are expressed as mean ± SEM. \**P<*0.05, \*\**P<*0.01, \*\*\**P<*0.001 in comparison to control.

### 3.3. Effects of Astaxanthin on free radical scavenging

ASX exerted the concentration-dependent inhibitory effect ranging from 7.75 ± 2.08 – 85.59 ± 2.04 % (*p<*0.05 - 0.001; 1.6 - 5000 µg/mL) on DPPH significantly (Table 1). On the other hand, the positive control, L-ascorbic acid, demonstrated increasing DPPH scavenging activity with accelerating doses (1.6 - 5000 µg/mL), exhibiting a 14.17 ± 2.04 - 97.08 ± 2.04 % (*p>*0.01 - 0.001; 1.6-5000 µg/mL) inhibitory effect on DPPH (Table 1).

**Table 1.**
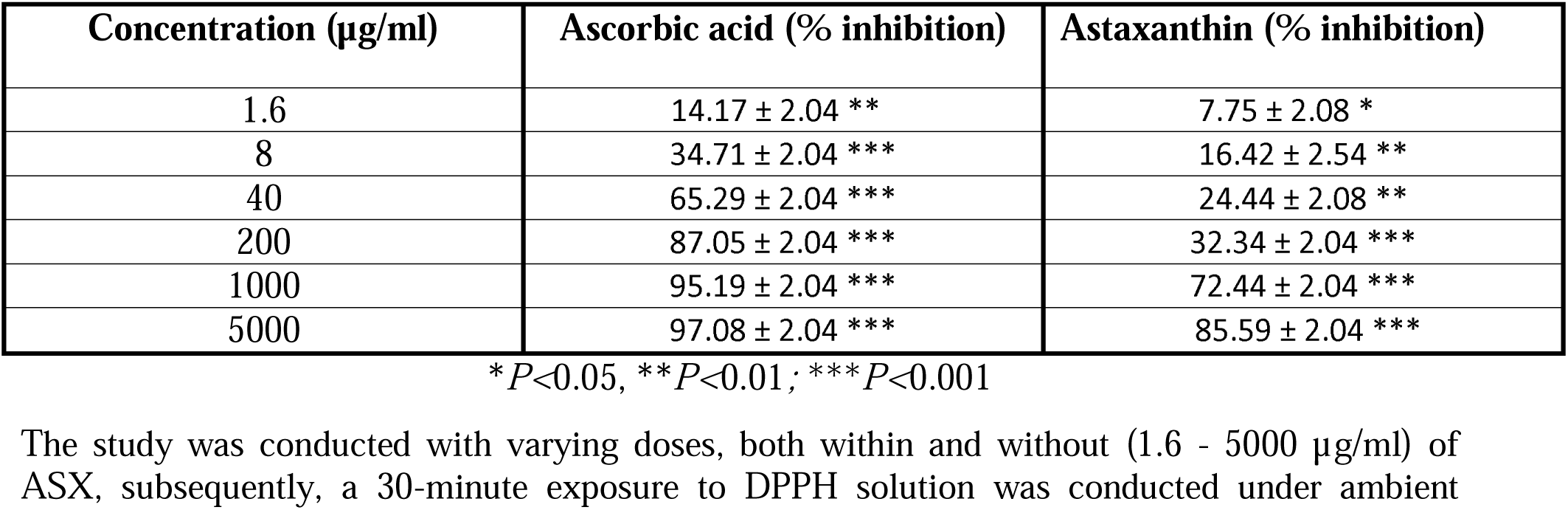
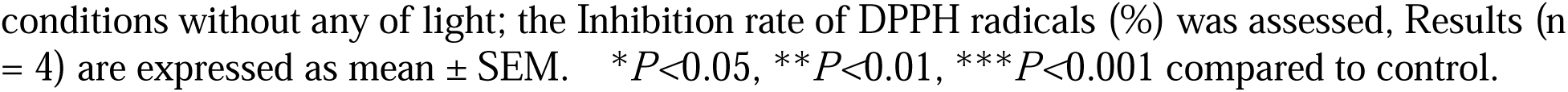
DPPH scavenging potential of L-ascorbic acid and ASX in a dose-dependent manner.

| Concentration (μg/ml) | Ascorbic acid (% inhibition) | Astaxanthin (% inhibition) |
| --- | --- | --- |
| 1.6 | $14.17 \pm 2.04$ ** | $7.75 \pm 2.08$ * |
| 8 | $34.71 \pm 2.04$ *** | $16.42 \pm 2.54$ ** |
| 40 | $65.29 \pm 2.04$ *** | $24.44 \pm 2.08$ ** |
| 200 | $87.05 \pm 2.04$ *** | $32.34 \pm 2.04$ *** |
| 1000 | $95.19 \pm 2.04$ *** | $72.44 \pm 2.04$ *** |
| 5000 | $97.08 \pm 2.04$ *** | $85.59 \pm 2.04$ *** |

\* $P$ <0.05, \*\* $P$ <0.01; \*\*\* $P$ <0.001
The study was conducted with varying doses, both within and without (1.6 - 5000 μg/ml) of ASX, subsequently, a 30-minute exposure to DPPH solution was conducted under ambient
conditions without any of light; the Inhibition rate of DPPH radicals (%) was assessed, Results (n = 4) are expressed as mean ± SEM. \* $P < 0.05$ , \*\* $P < 0.01$ , \*\*\* $P < 0.001$ compared to control.

### 3.4. Effects of Astaxanthin on Metabolic Study

ASX (30 and 60 mg/5 ml/kg b.w.) significantly decreased (*p<* 0.01) at 12 hours nocturnal meals and post-24-hour intake (p < 0.001) compared to HFF-induced type 2 diabetic mice (Figure 3A). Additionally, ASX (30 and 60 mg/5 ml/kg b.w.) improved fluid intake, as well as stool and urine output, were monitored during the evening hours between 6:00 and 8:00 PM though was not significant compared to HFF-induced type 2 diabetic mice (Figure 3B, D, E). However, ASX, at (60 mg/5 ml/kg, b.w.) was more effective (*p<*0.001) than at 30 mg/5 ml/kg, b.w. (Figure 3A). The positive control, metformin also significantly decreased food intake at 12- and 24-hours’ time interval (*p<*0.001; Figure 3A) and also improved fluid intake, as well as stool and urine output, were monitored during the evening hours between 6:00 and 8:00 PM though were not significant compare to HFF control (Figure 3B, D, E).

**Figure 3:**
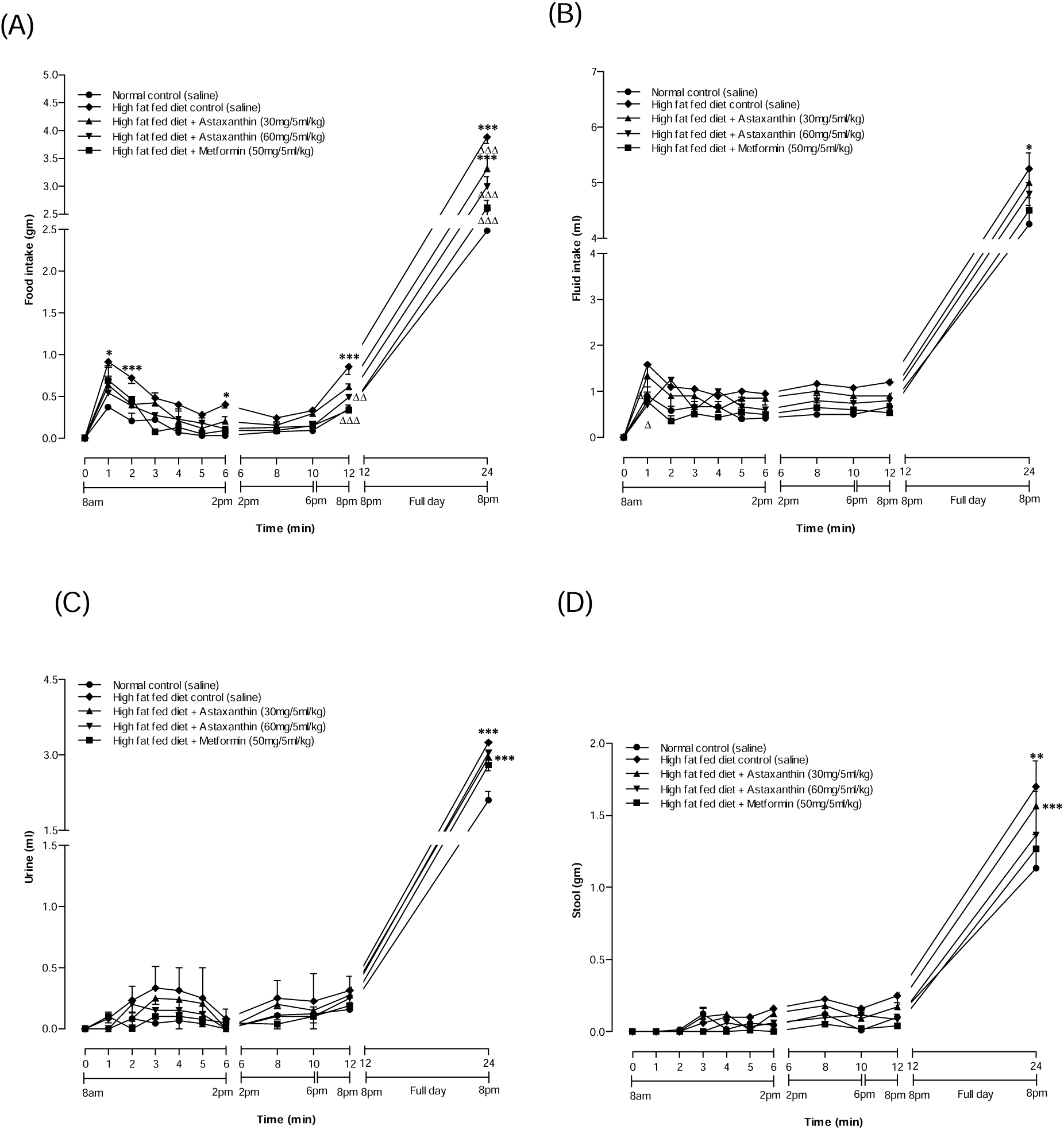
Results of Astaxanthin on metabolic parameters (A) food intake (B) Fluid intake (C) Urine and (D) Stool in HFF mice. The parameters were noted at 1, 2, 3, 4, 5, 6, 8, 12 and 24 hours. n = 6. Results are presented as mean ± SEM. **\****P<*0.05**, \*\****P<*0.01**, \*\*\****P<*0.001 compared to control and ^Δ^*P<*0.05, ^ΔΔ^*P<*0.01, ^ΔΔΔ^*P<*0.001 compared to HFF induced type 2 diabetic mice.

### 3.5. Effects of Astaxanthin on acute oral glucose tolerance test

Oral gavage of ASX at (60 mg/5 ml/kg, b.w.), along with glucose (18 mmol/kg, b.w.) to the respective groups exhibited in a significant enhancement in oral glucose tolerance at 30 and 60 minutes on days 15 and 30, and at 30 and 120 minutes on day 0 in HFF type 2 diabetic mice (*p<*0.05 - 0.001; Figure 4), in comparison to the HFF diabetic control group. Among the three tests, ASX at 60 mg/5 ml/kg showed improvements in glucose tolerance with most significant decrease in blood glucose at 15 and 30 days (*p<*0.001; Figure 4B, 4C) in 30 and 60-mins time points. The positive control, metformin also significantly improved in oral glucose tolerance at 30 and 60 mins on days 0,15 and 30 in HFF type 2 diabetic mice (*p<*0.001; Figure 4), compared to the HFF diabetic control group.

**Figure 4:**
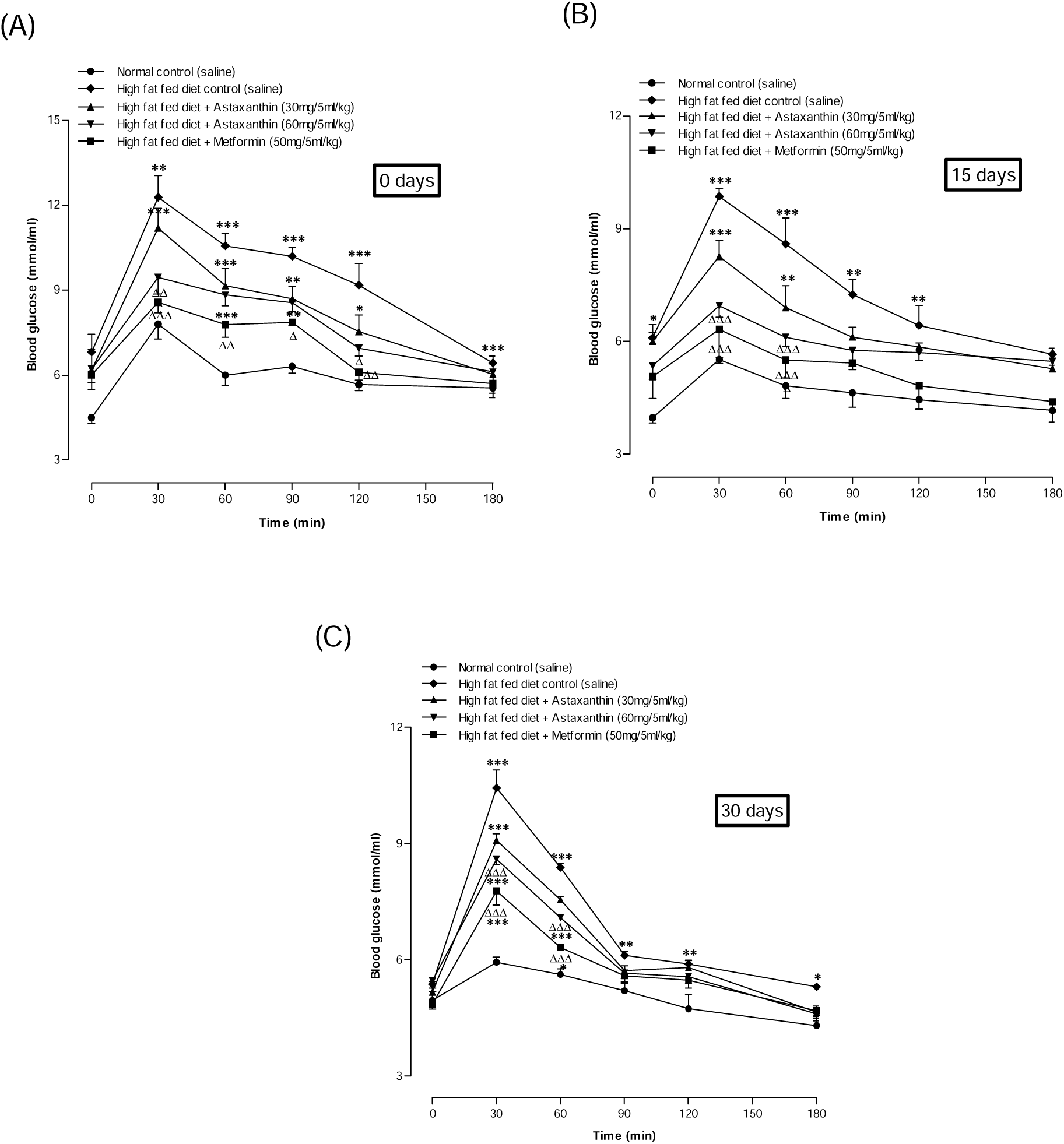
Effects of Astaxanthin on oral glucose tolerance at (A) 0 days, (B) 15 days and (C) 30 days. Tests were performed following administration of ASX (30 and 60 mg/5 ml/kg, b.w.) and glucose (18 mmol/kg, b.w.) in HFF induced type 2 diabetic mice. The Blood glucose levels were evaluated at 0, 30, 60, 90, 120 and 180-min. n = 10. Results are expressed as mean ± SEM. **\****P<*0.05**, \*\****P<*0.01**, \*\*\****P<*0.001 compared to control and ^Δ^*P<*0.05, ^ΔΔ^*P<*0.01, ^ΔΔΔ^*P<*0.001 compared to high fat-induced type 2 diabetic mice.

### 3.6. Chronic effects of astaxanthin on blood glucose, body weight, food, and fluid intake

twice daily oral gavage of ASX at two different concentrations (30 and 60 mg/5 ml/kg b.w.) in HFF induced diabetic mice exhibited a significant and consistent lowering fasting blood glucose readings. ASX at (30 mg/5 ml/kg b.w.) exhibited results from 15 to 33 days, whereas ASX at 60 mg/5 ml/kg demonstrated improvements from 9 to 33 days (*p<*0.01 - 0.001; Figure 5A) compared HFF control. On the other hand, ASX at (60 mg/5 ml/kg b.w.) exhibited the reduction of body weight from day 24 to day 33 (*p<*0.05 - 0.001; Figure 5B) in comparison to diabetic control. Finally, both the food and fluid consumption at ASX 60 mg/5 ml/kg group showed the most significant reduction than all groups and the reduction started observable from day 24 and 21 to 33 (*p<*0.01 - 0.001; Figure 5C, D), respectively. The positive control, metformin also significantly improved the chronic fasting blood glucose, body weight and frequency of food and fluid intake (*p<*0.05 - 0.001; Figure 5B, D, E).

**Figure 5:**
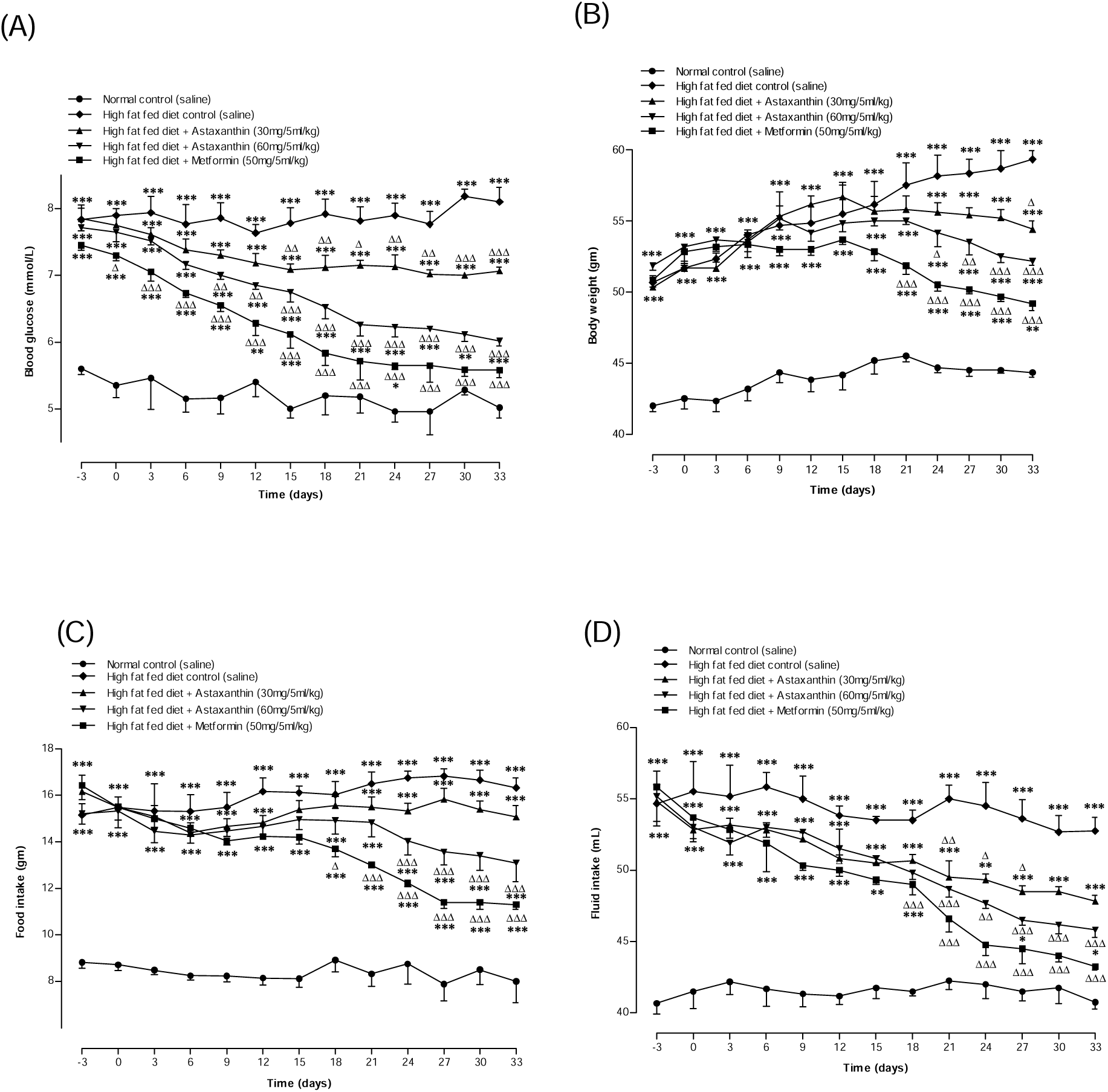
Chronic impacts of Astaxanthin 30 and 60 mg/5ml/kg on (A) blood glucose, (B)body weight, (C) food and (D) fluid intake. Tests were conducted after a twice-a-day oral dose of ASX (30 and 60 mg/5 ml/kg, b.w.) and metformin (50 mg/5 ml/kg b.w.) in HFF induced type 2 diabetic mice. Results (n = 6) are expressed as mean ± SEM. **\****P<*0.05**, \*\****P<*0.01**, \*\*\****P<*0.001 compared to control and ^Δ^*P<*0.05, ^ΔΔ^*P<*0.01, ^ΔΔΔ^*P<*0.001 compared to HFF induced type 2 diabetic mice.

### 3.7. Effects of Astaxanthin on Gastrointestinal motility

The ingestion of ASX at the concentration of (30 and 60 mg/5 ml/kg b.w.) showed a notable enhancement in gastrointestinal motility (*p<*0.05; Figure 7A). Correspondingly, the positive control metformin (50 mg/5 ml/kg b.w.) also showed significance increased (*p<*0.01; Figure 7A).

### 3.8. Effects of Astaxanthin on unabsorbed sucrose content in the gut

A significant increase (*P<*0.05 and *P<*0.01; Figure 6A, B, C) was observed unabsorbed sucrose content at 30 and 60 minutes in the stomach, upper, and middle segments of the small intestine subsequent oral dosing of sucrose (2.5 g/kg b.w.) and the ASX (30 and 60 mg/5 ml/kg b.w.). Additionally, a significant rise was noted across the cecum and large intestine after two hours (*P<*0.05; Figure 6E, F). At the 4-hour interval, significant amount of sucrose was still detectable in the large intestine of the ASX (60 mg/5 ml/kg b.w.) treated group (*P<*0.05; Figure 6F).

**Figure 6:**
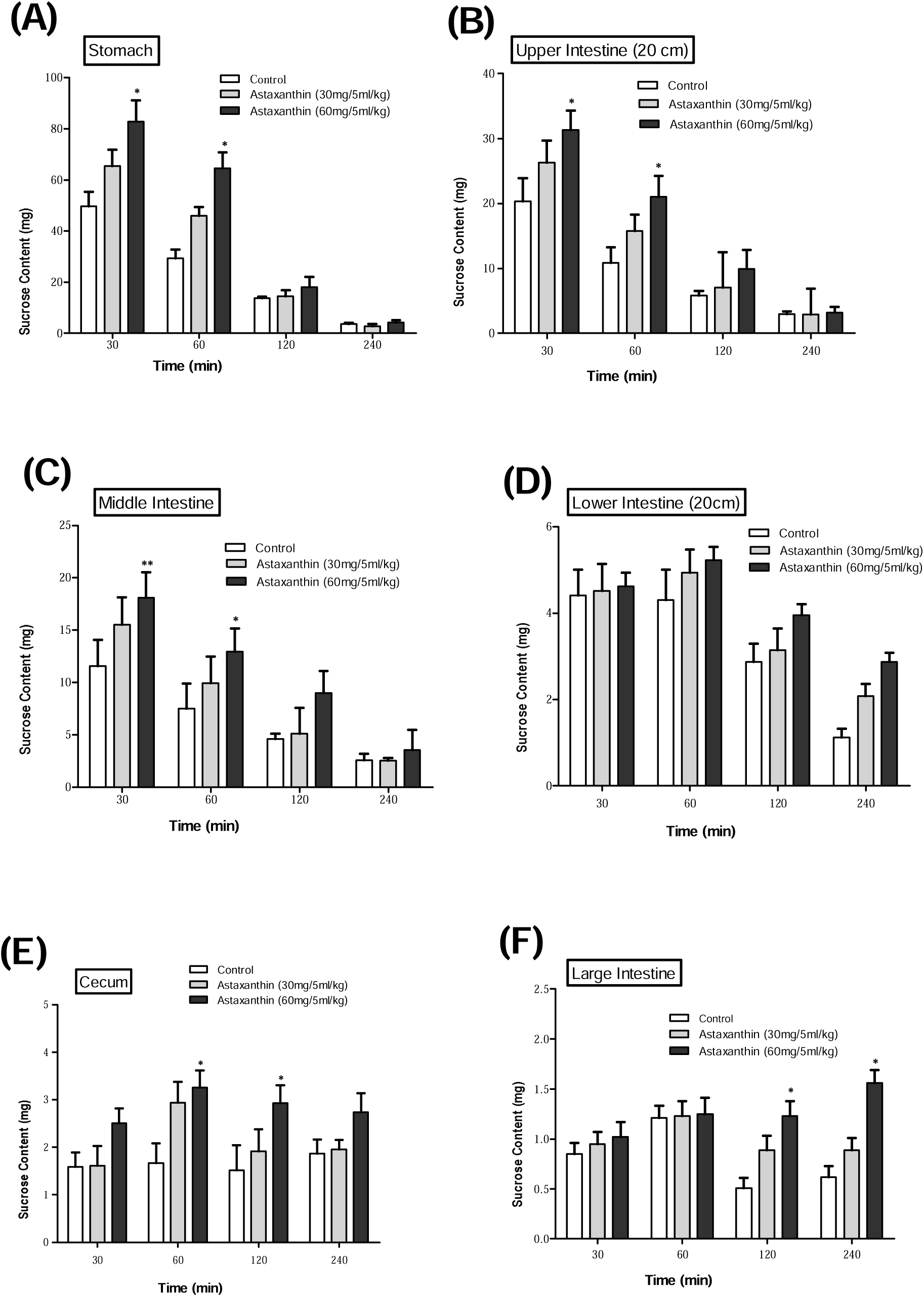
Gastrointestinal residual unabsorbed sucrose content in (A) Stomach, (B) Upper Intestine, (C) Middle Intestine, (D) Lower Intestine, (E) Cecum, and (F) Large Intestine. Type 2 diabetic rats were fasted for 24 hours prior to receiving an oral sucrose solution (2.5 g/kg b.w.), either alone (control group) or in combination with *ASX* (30 and 60 *mg/5 ml/kg b.w.*). Results (n = 6) are expressed as mean ± SEM. \**P*<0.05, \*\**P*<0.01, \*\*\**P*<0.001 compared to control.

### 3.9. Effects of Astaxanthin on liver glycogen and lipid profile

Following 33 days of oral administration of ASX at (30 and 60 mg/5 ml/kg, b.w.) alone with ASX (60 mg/5 ml/kg b.w.) show a significant (*p<*0.01) increase in the glycogen content of the liver was noted and metformin as a positive control also significantly (*p<*0.001) increase liver glycogen content (Figure 7B). ASX showing a significant improvement in the blood plasma cholesterol assessment characterized by a reduction in systemic triglycerides (*P<*0.05; Fig. 7C), total cholesterol (*P<*0.01; Figure 7D), LDL (*P<*0.01; Figure 7E), and VLDL cholesterol contents (*P<*0.001; Figure 7F) in comparison to the HFF induced hyperlipidemic mice.

## 4. Discussion

Diabetes is an alarming global threat to early mortality [40]. Current treatments are costly, unable to prevent serious complications of diabetes, and are not widely available [41]. Previous research suggests that administering Astaxanthin (ASX), whether orally or by injection, enhances insulin sensitivity, decreases high blood sugar levels, and protects against retinopathy, nephropathy, and neuropathy [42]. Although previous studies have investigated the antihyperglycemic action of ASX, there is insufficient suppression of carbohydrate digestion and absorption and GI motility. The present research investigated the antidiabetic effects of ASX and its mechanisms using *in vitro* starch digestion, glucose diffusion, free radical scavenging and *in vivo* experiments, including OGTT, gastrointestinal motility, liver glycogen content and lipid profile tests in hyperlipidemic albino mice. Acute and chronic metabolic effects were assessed through observations of food and water intake, urination, and defecation in high-fat-fed albino mice.

An effective strategy for managing post-prandial blood sugar levels is the suppression of the digestive enzymes, α-amylase and α-glucosidase [43]. In our study, *In-vitro* starch digestion was performed to examine the impact of ASX on the activity of α-amylase and α-glucosidase enzymes in the release of glucose from carbohydrate digestion. The results revealed a significant reduction of enzymatic digestion of starch on ASX. This finding aligns with the known inhibitory effects of Acarbose, the positive control in our experiment, which also displayed a concentration-dependent inhibition of starch digestion [44]. Recent studies demonstrate that ASX markedly reduces enzymatic glucose digestion, revealing its ability to inhibit starch digestion and hence efficiently regulate blood glucose levels [45][46].

The reduction of gastrointestinal absorption and glucose diffusion is an important strategy for regulating the glycemic response to meals [47]. In additionally, glucose diffusion assay is an effective in vitro technique for predicting the impact of fiber on the delay of glucose absorption in the gastrointestinal tract. [48]. In this study, the impact of ASX on glucose transport was examined utilizing a basic in vitro dialysis-based model. These findings correspond with prior research indicating that astaxanthin could affect glucose absorption [49], [50].

Recent study indicates that diabetes is correlated with oxidative stress and the formation of free radicals, thereby resulting in insulin resistance [51]. This *in vitro* DPPH study revealed the significant antioxidant potential of ASX, demonstrating Its scavenging capacity for free radicals thus addressing oxidative stress, a key factor in diabetes and its complications. Previous research showed that ASX was reported to reduce hyperglycemia-induced oxidative damage in β-cells, enhancing glucose tolerance, elevating insulin levels, and considerably lowering blood glucose levels [52].

Excessive amount meals increase the chance of overweight and insulin resistance, and T2DM [53]. Metabolic studies were conducted to examine the impact of ASX on various variables, such as food and fluid intake, along with stool and urine production in HFF-induced type 2 diabetic Swiss albino mice. ASX was observed to decrease all four aforementioned parameters, particularly during night time. In the previous research, on high-fat-fed mice with type 2 diabetes, ASX therapy showed reduced amount of food and fluid intake, urine and feces formation after 24 hours [54]

Effective management of postprandial hyperglycemia is essential for mitigating and preventing adverse reactions correlated with type 2 diabetes [55]. In this acute OGTT, ASX along with glucose showed ameliorative effects in HFF type 2 diabetic mice. Particularly, the effectiveness of ASX at high concentrations (60 mg/5 ml/kg, b.w.) Produced a markedly significant hypoglycemic effect at the 30 and 60 minutes in the 15 and 30-day time. In the previous research, involving prediabetic patients demonstrated lower blood glucose and glycated hemoglobin readings following an OGTT conducted over a period of 120 minutes [56]. Although numerous results indicate that oral treatment of ASX dramatically reduced pre-prandial glycemic levels in db/db mice, a recognized design of obesity for T2DM [57], [58], [59].

In this present study, after taking ASX orally twice a day for 33 days, we noticed a considerably lower body weight, food and fluid intake, and fasting blood glucose. In the previous research, ASX treated with 6 mg/kg for 45 consecutive days considerably lowered blood sugar and insulin responses in mice administered a higher fructose and fat-rich diet [60]. In addition, the prolonged medication of ASX at a concentration of 35 mg/kg over a period of 12 weeks found a Glycemic control effect [61]. Furthermore, previous report also showed that after 60 days of oral ASX therapy, high-fat-fed type 2 diabetic mice showed reduced alimentary consumption body mass [54].

The decrease in transit time in the gastrointestinal tract leads to a lack of duration for the digestion and absorption of glucose [62]. Gastrointestinal motility has been monitored using a solution of BaSO4 milk. The present study indicates that ASX improves gut motility, claiming a potential reduction in the time span of carbohydrate digestion and absorption in the gastrointestinal tract, which leads to less glucose absorption [41], [63].

The noted reduction in sucrose content may be ascribed to the suppression of enzymatic function in the gastrointestinal tract [64]. The current investigation revealed that ASX can impede sucrose absorption across various segments of the gastrointestinal system, indicated by significant quantities of unabsorbed sucrose remaining in the gut. The effects of ASX are comparable to acarbose, an α-glucosidase inhibitor recognized for its efficacy in diminishing post-meal hyperglycemia in individuals with diabetes [65].

This study showed an increase in liver glycogen levels, indicating pancreatic beta-cell regeneration and reduced fat deposition [66]. The lipid profile assessments of the ASX revealed a substantial decrease in blood serum triglycerides, total cholesterol, low-density lipoprotein, and very low-density lipoprotein levels. The present results align with prior research on ASX, which shown a decrease in liver weight, liver triglyceride, plasma triglyceride, and total cholesterol [67]. Additionally, it has been linked to significant Impact on various lipid parameters, such as decreased total cholesterol, LDL, and triglyceride levels, as well as diminished HDL [68], [69].

## 5. Conclusion

The present study reveals that ASX inhibits carbohydrate digestion and absorption, exerts inhibitory effects on DPPH and increases gastrointestinal motility. Thus, current findings establish the role of ASX as a useful dietary supplement for managing diabetes mellitus and related complications. Further research involving ASX is needed to explore new therapeutic strategies against diabetes and associated disorders.

## Abbreviations

ASX – Astaxanthin, DM - Diabetes Mellitus, T1DM - Type 1 Diabetes Mellitus, T2DM - Type 2 Diabetes Mellitus, MODY - Maturity Onset Diabetes of the Young, b.w. – Body weight, OGTT - Oral Glucose Tolerance Tests, GI – Gastrointestinal, DPPH - 2,2-Diphenyl-1-picrylhydrazyl, ICDDRB - International Center for Diarrheal Disease Research, Bangladesh, HFF - High-fat feed, GOD-PAP - Glucose Oxidase/four-Aminophenazone-Phenol, CHOD-PAP - Cholesterol Oxidase-Peroxidase, GPO-PAP - Glycerol Phosphate Oxidase-Peroxidase.

## Author Contributions

P.A. was accountable for the conceptualization and structure of this research and collaboratively supervised its execution; M.S.R., P.A., and J.A.S. contributed to the conducted experiments and data analysis; M.S.R. undertook the interpretation of these results, figure preparation, and primary manuscript drafting; and M.S.R. and J.M.A.H. participated in editing the revised manuscript. The contributors have reviewed and consented to the final draft of this writing.

## Funding

This study acquired no additional support.

## Acknowledgments

The authors appreciate the Independent University, Bangladesh (IUB), Dhaka, for providing the facilities and resources that facilitated the performance of this study.

## Conflict of Interest

The authors claim that there is no disagreement of obligation.

## Notes

### Competing Interest Statement

The authors have declared no competing interest.

